# Genetic mapping of craniofacial traits in the Mexican tetra reveals loci associated with bite differences between cave and surface fish

**DOI:** 10.1101/2022.11.23.517717

**Authors:** Amanda K. Powers, Carole Hyacinthe, Misty Riddle, Young Kwang Kim, Alleigh Amaismeier, Kathryn Thiel, Brian Martineau, Emma Ferrante, Rachel Moran, Suzanne McGaugh, Tyler Boggs, Joshua B. Gross, Clifford J. Tabin

**Affiliations:** Department of Genetics, Blavatnik Institute at Harvard Medical School, 77 Avenue Louis Pasteur, Boston, MA 02115; Department of Biology, University of Nevada, Reno, 1664 N. Virginia St., Reno, NV 89557; Harvard School of Dental Medicine, 188 Longwood Ave., Boston, MA 02115; Department of Biology, Xavier University, 3800 Victory Pkwy., Cincinnati, OH 45207; Department of Biology, Texas A & M University, 100 Butler Hall, College Station, TX 77843; Department of Ecology, Evolution and Behavior, University of Minnesota, 1500 Gortner Ave., Saint Paul, MN 55108; Department of Biological Sciences, University of Cincinnati, 312 College Dr., Cincinnati, OH 45221

**Keywords:** Jaws, teeth, Meckel’s cartilage, Quantitative trait loci, cavefish

## Abstract

The Mexican tetra, *Astyanax mexicanus*, includes interfertile surface-dwelling and cave-dwelling morphs, enabling powerful studies aimed at uncovering genes involved in the evolution of cave-associated traits. Compared to surface fish, cavefish harbor several extreme traits within their skull, such as a protruding lower jaw, a wider gape, and an increase in tooth number. These features are highly variable between individual cavefish and even across different cavefish populations. To investigate these traits, we created a novel feeding behavior assay wherein bite impressions could be obtained. We determined that fish with an underbite leave larger bite impressions with an increase in the number of tooth marks. Capitalizing on the ability to produce hybrids from surface and cavefish crosses, we investigated genes underlying these segregating orofacial traits by performing Quantitative Trait Loci (QTL) analysis with F_2_ hybrids. We discovered significant QTL for bite (underbite vs. overbite) that mapped to a single region of the *Astyanax* genome. This work highlights cavefish as a valuable genetic model for orofacial patterning and will provide insight into the genetic regulators of jaw and tooth development.

## Introduction

One of the hallmarks of early vertebrate evolution is the biting jaw (de Beer, 1937; Romer, 1941). Because the mandibular arch can be found in jawless fishes such as lamprey and hagfish, it is likely that the morphological identity of lower jaw components (i.e. pharyngeal arches) was present in a common ancestor to the jawless cyclostomes and jawed gnathostomes (Kuratani, 2012). Among other cranial bones, the lower jaw is highly conservative across vertebrates from extinct armored placoderms to living tetrapods (Long, 2016), suggesting conserved genetic networks govern jaw development.

The emergence of diversity in jaw morphology is linked to feeding ecology (Hill et al. 2018). Classic examples of adaptive radiations, such as beak shape in Darwin’s finches (Abzhanov et al. 2006) and jaw diversity in cichlids (Husley et al. 2010), occur through the expansion into new feeding niches, leading to extreme changes in morphology and in some cases speciation events. Cichlids exhibit a spectrum of variation in their oral jaws, from short jaws amenable to biting hard surfaces to elongated jaws for suction feeding (Powder & Alberston, 2016). The emergence of these morphological changes is integrated in environmental and ecological pressures.

Perhaps one of the most extreme environmental pressures an organism can face is the subterranean habitat. Obligate cave-dwellers face perpetual darkness, scarce food sources and isolation from other ecosystems. Despite these challenges, cave organisms thrive in this environment. For example, *Astyanax mexicanus* cavefish have evolved physiological and morphological traits suited for life in complete darkness, such as starvation resistance (Aspiras et al. 2015; Xiong et al. 2018, Riddle & Aspiras et al. 2018), enhancement of sensory systems (Jeffery, 2001; Yoshizawa et al. 2014; Wilkens, 2020), sleep loss/constant foraging (Duboué et al. 2011), and changes to their immune system (Peuß et al. 2020), relative to extant surface-dwelling fish. In addition to these changes, cavefish harbor extreme changes in morphology, including several craniofacial traits, such as cranial bone fragmentations and spontaneous fusions, as well as fluctuating and directional asymmetries (Gross & Powers, 2020). These craniofacial features are highly variable across both individual cavefish, as well as the ∼30 known *Astyanax* cavefish populations found in northeastern Mexico. Within their oral jaws, adult cavefish exhibit an increase in both upper and lower jaw dentition (tooth number) compared to surface fish (Atukorala et al. 2013). Further, larval cavefish have wider and more protruding lower jaws (Jeffery, 2001; Yamamoto et al. 2009).

An elongation of the lower jaw is not unique to the blind Mexican cavefish, however. Protruding lower jaws have been characterized in cavefish across the globe including the Chinese cavefish (*Sinocyclocheilus*; Ma et al. 2019), the cavefish of the Ozarks (*Amblyopsis rosae*; Romero, 2009), and an Australian cavefish (*Milyeringa brooksi*; Chakrabarty, 2010). This parallel evolution of changes in the lower jaw suggests a possible adaptive significance.

Toward that end, we set out to characterize changes in lower jaw morphology in adult *Astyanax* cavefish using morphological, behavioral, and genetic analyses. We discovered that the wider, protruding lower jaws observed in larval cavefish persist in the adult cranium, resulting in an underbite compared to the slight overbite or normal occlusion found in surface fish. To determine if the underbite is of functional importance, we assessed the maximum gape (mouth opening) and feeding behavior using a novel feeding assay. Further, we capitalized on the ability to generate viable hybrids from surface x cavefish crosses and employed a genetic association study to illustrate that bite differences are under genetic control in *A. mexicanus*. Next, we were able to pinpoint an associated region in the genome and generate a subsequent list of candidate genes for this trait. Together, our analyses reveal a novel role for differences in jaw morphology and tooth patterning in cavefish that likely evolved as an alternative feeding strategy in nutrient poor caves.

## Materials and Methods

### Fish Husbandry and Specimens

Fish were bred and maintained in the laboratory of Dr. Clifford Tabin at Harvard Medical School on a custom recirculating system (Temperature: 23C, pH: 7-7.5, and Conductivity: 1200-1400uS) under a 12 hour light/dark cycle. F_1_ hybrids were generated from a paired mating of male *Astyanax mexicanus* surface fish (derived from the Río Choy river) and female Pachón cavefish. The genetic mapping pedigree was made up of F_2_ hybrids (n=219) from three clutches from F_1_ surface x Pachón hybrid siblings. For behavioral analysis a second F_2_ population (n=30) was generated from a single cross of F_1_ siblings. It has been previously determined that there is no maternal effect on jaw morphology for hybrid crosses by looking at reciprocal hybrids (Ma et al. 2018). All procedures were approved under IACUC protocol (#IS00001612).

### Feeding behavior assay

We created “food carpet” molds that were placed at the bottom of assay tanks, from which we could recover bite impressions. Solidified gelatinous food carpets were made using comestible gelatin (Knox). Gelatin powder was melted in boiled, filtered reverse osmosis water and mixed with a solution base of infused fish pellets (New Life Spectrum Thera+A) using an electric kettle (Muller) in a ratio of 1:1. The warm liquid mix was poured into silicone molds, chilled at room temperature and stored overnight at 4°C for solidification. Molds with solidified gelatin were placed at the bottom of recording tanks filled with water, occupying the entire bottom of the tank as a “food carpet” (Fig. 2A-D).

The feeding behavior assay was performed on n=5 surface fish, n= 5 Pachón cavefish as well as n=20 F_2_ hybrids recorded in 1.7L tanks, set up in an insulated chamber. Each fish was recorded in complete dark conditions for 1h with a HD infrared camera (Grundig Pro, Germany). All assays were watched live to control for actual feeding episodes; feeding episodes are described here as active mouth-picking on the “food carpet”. After 1h of trial, fish were returned to their housing tank and food carpets were extracted from recording tanks. Food carpets were dried for ∼12 hrs in a low humidity room and imaged under a light stereomicroscope (Leica M165FC) at 32x magnification (Fig. 2E-H).

Tanks were filmed via a front-facing camera and videos were acquired through Open Broadcaster Software (OBS) studio in “.avi” format. Videos were manually analyzed with Odrec software (S. Pean, IFREMER, France) to quantify the average and maximum body angles adopted by the fish over 1h periods for each feeding episode (Fig. S1). Accurate measures of the body angle were facilitated with a protractor overlayed directly on the tank in 10° quadrants (Fig. 2A-D).

### Phenotypic analysis

To assess the maximum mouth opening (“gape”), specimens (n=8 from each group) were sacrificed using a lethal dose (400ppm) of tricaine (MS-222; Sigma) and immediately imaged under light microscopy at 20x before rigor mortis set in to maintain flexibility in the jaw joints. Upper and lower jaws were pinned using Styrofoam backing at the maximum gape (Fig. 1A, D). Gape was measured as the angle at the intersect of the maxillary and dentary (lower jaw) bones using the angle tool in ImageJ software (v2.0.0-rc-69). An analysis of variance (ANOVA) and post hoc Tukey’s HSD were performed using R studio software (v2022.07.2; Table S1). Lower jaw length was measured in F_2_ hybrids (n=186) using the line tool in ImageJ and normalized to fish standard length (Fig. 2I). For pairwise comparisons, a t-test comparison of means (StatPlus:mac LE v6.2.21) was used to test for statistical significance.

**Figure 1.**
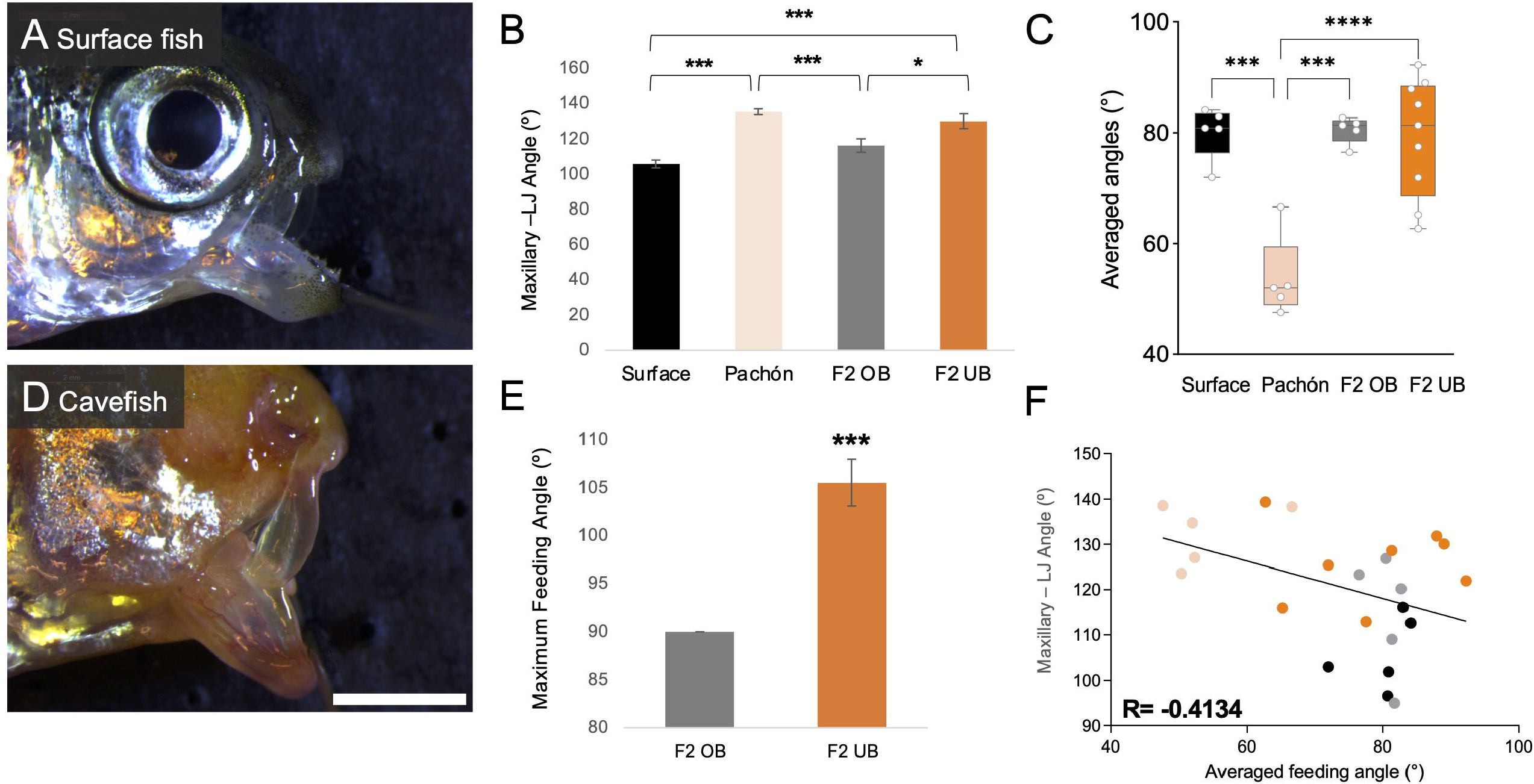
Fish with an underbite exhibit a larger maxillary – lower jaw angle (gape), which negatively correlates with feeding angle. Adult surface fish (A) have a smaller maxillary – lower jaw angle (mean= 106°) compared to cavefish (D; mean= 136°; p<0.001) (B). F_2_ hybrids scored as having an overbite were not significantly different than surface fish (mean= 116.5°; p=0.1) (B). Additionally, F_2_ hybrids scored as having an underbite were not significantly different from cavefish (mean= 130°; p=0.62) (B). In agreement with data from Kowalko et al. 2013, we determined that surface fish feed at an average of ∼80-90° angle compared to cavefish that feed at ∼45° (C). While F_2_ hybrids with an overbite feed at a similar angle to surface fish, F_2_s with an underbite feed within a wide averaged range between 65-95° (C). Compared to F_2_s with an overbite that have a maximum feeding angle of 90°, underbite F_2_s had a significantly higher maximum feeding angle at 110° (E). There is a negative correlation (R= -0.4134) between a higher maxillary – lower jaw angle and feeding posture angle (F). White scale bar set at 2 mm.

**Figure 2.**
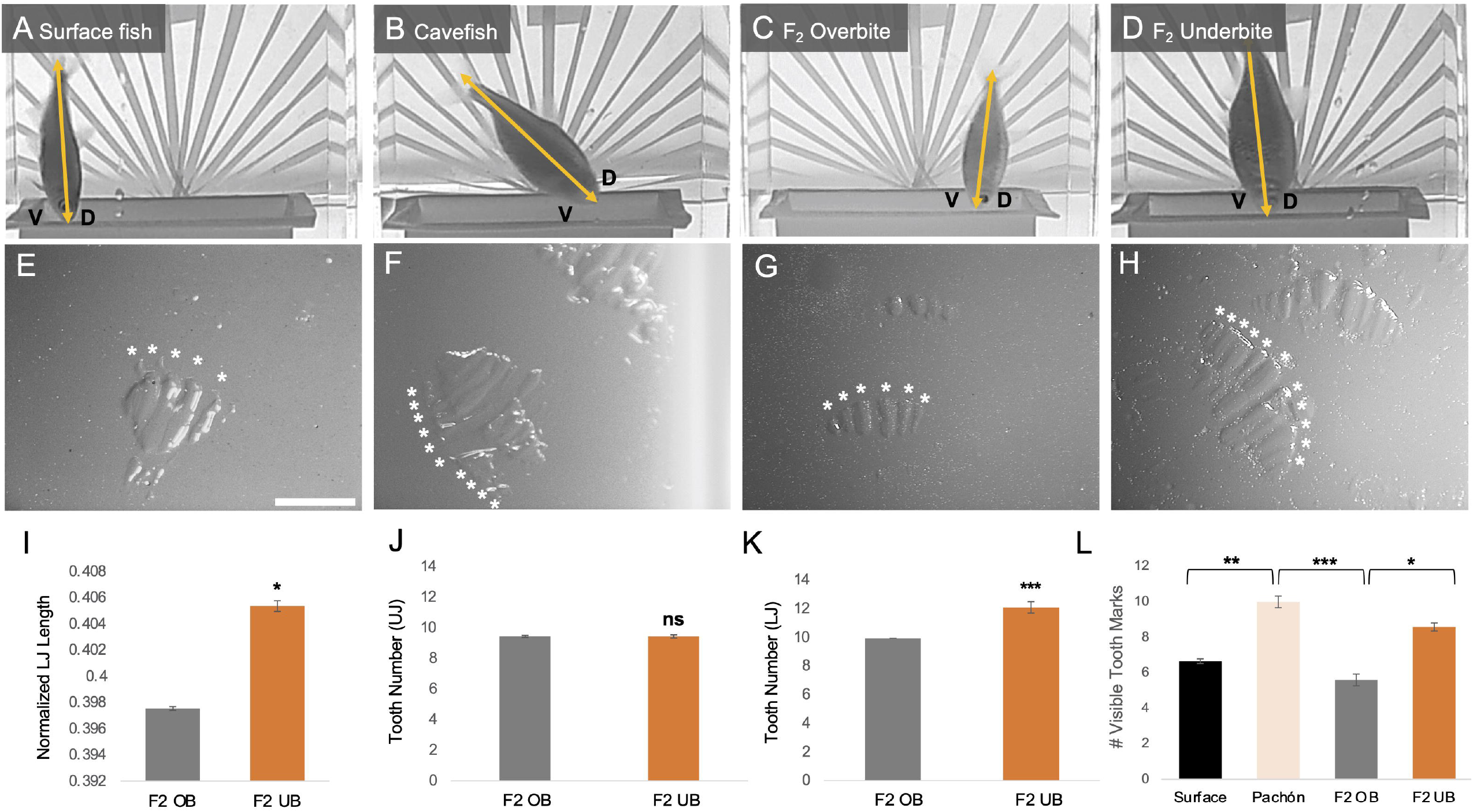
Fish with an underbite leave a greater number of tooth marks in bite impressions compared to fish with an overbite. Surface fish (A, E) make bite impressions with fewer tooth marks (mean= 6.7) compared to cavefish (mean= 10; p<0.01) (B, F, L). Accordingly, F_2_ hybrids with an overbite (C, G) left fewer tooth marks (mean= 5.6) than F_2_ hybrids with an underbite (mean= 8.6; p<0.05) (D, H, L). F_2_ hybrids with an underbite have significantly longer lower jaws (normalized length; p<0.05) (I) and an increase in lower jaw tooth number (p<0.001) compared to F_2_ hybrids with an overbite (K). There was no significant difference between upper jaw tooth number between the two hybrid groups (J). White scale bar set at 1 mm.

For three-dimensional analysis, high resolution micro-computed tomography (MicroCT) imaging was performed at the Center for Advanced Orthopaedic Studies at the Beth Israel Deaconess Medical Center (Boston, MA). MicroCT scans were performed on Pachón cavefish (n=5), surface fish (n=5), F_1_ (n=5) and F_2_ hybrids (n=219) at 15uM resolution producing ∼500 DICOM formatted images per specimen that were reconstructed into a single three-dimensional volume rendered file using Amira software (v6.0; FEI Company, Hillsboro, OR) according to methods outlined in Powers et al. 2017.

### Quantitative Trait Loci (QTL) analysis

R/qtl (v1.46-2; Broman et al. 2003) was used to perform QTL analysis according to methods outlined in Riddle et al. 2020. Briefly, a linkage map was constructed using loci identified from genotyping-by-sequencing (GBS) technology. The linkage map consisted of 1,839 GBS markers from 219 F_2_ individuals assembled into 25 linkage groups (Riddle et al. 2020). A genome-wide logarithm of odds (LOD) score was calculated for the “bite” phenotype. Bite was scored as a binary trait; overbite was scored as 0 and underbite was scored as 1 (Fig. 3A). Peak markers rising above the significant LOD threshold were extracted and phenotypic effect plots were generated to determine which genotypes were associated with bite differences (Fig. 3E).

**Figure 3.**
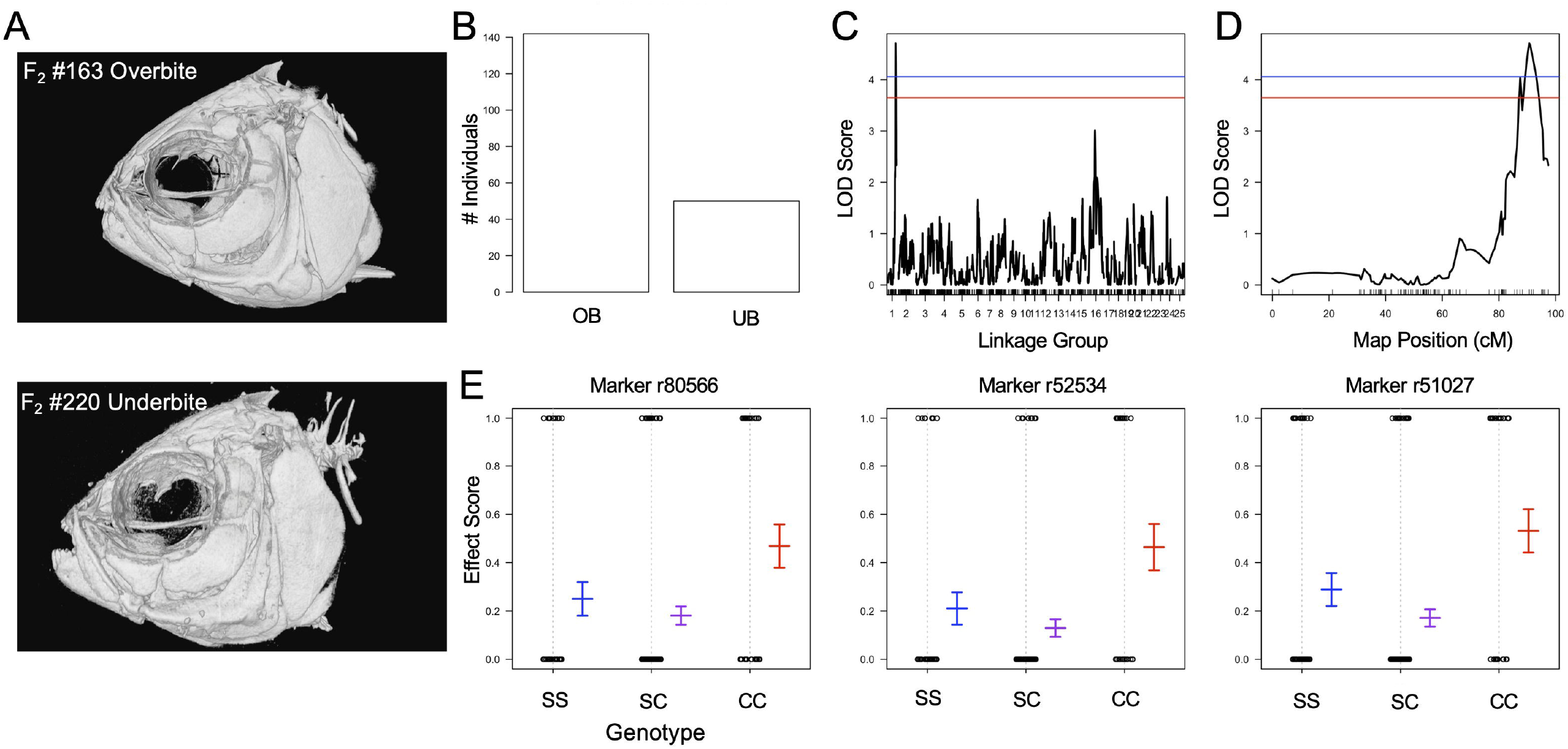
Quantitative Trait Loci (QTL) analysis reveals a genetic basis for bite differences. Representative F_2_ hybrid microCT images demonstrate bite differences scored as a binary trait for overbite (#163) and underbite (#220) (A). The frequency of F_2_ individuals exhibiting an overbite was ∼75%, while ∼25% of pedigree was scored as having an underbite (B). A single QTL peak was recorded for the bite phenotype rising about the significance threshold (blue line p<0.05; red line p<0.1) (C). The QTL peak resides on linkage group 1 between map positions 86-95 cM (D). Genetic marker r52534 had the peak LOD score (4.708) and the effect plot indicates that the homozygous cavefish genotype is associated with an underbite, while the homozygous surface fish and heterozygous genotypes are associated with an overbite (E). Flanking genetic markers r80566 and r51027 illustrate the same phenotypic effect (E).

Markers within the critical QTL region were mapped to the Pachón cavefish genome (AstMex102) scaffolds (Riddle et al. 2020) and the surface fish genome (*A. mexicanus* genome 2.0) chromosomes using the BLAST algorithm (Ensembl v108). The associated regions between the linkage map (LG1), cavefish genome scaffolds, and surface fish genome were visualized by generating a Circos plot (Fig. 4; Krzywinski et al. 2009). A candidate gene list was extracted from an ∼8Mb region on chromosome 7 using a custom pipeline outlined in Moran et al. 2022.

**Figure 4.**
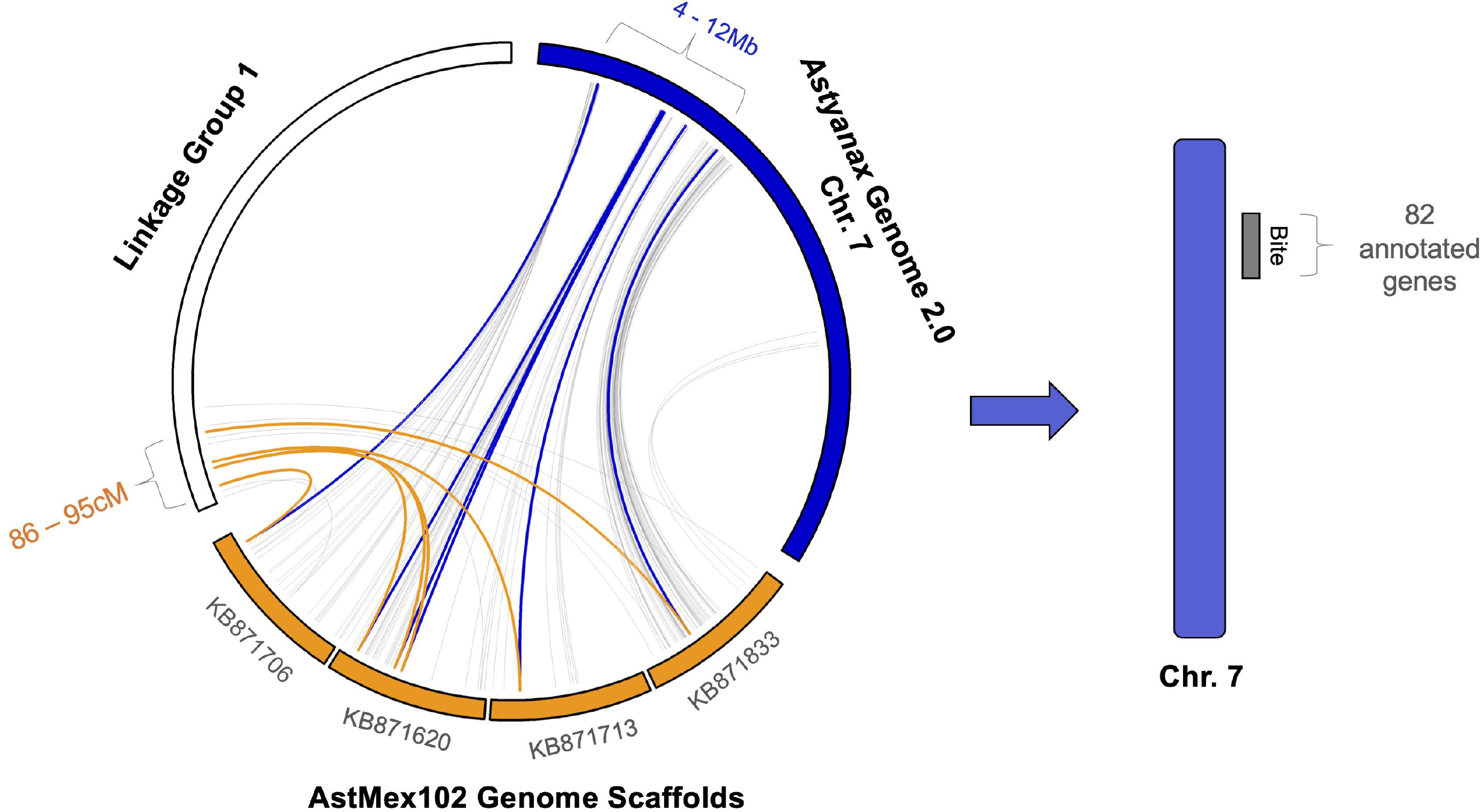
The peak QTL region maps to both the Pachón cavefish and surface fish genomes. Six genetic markers on linkage group 1 (86-95 cM) were anchored to four Pachón cavefish genome (AstMex102) scaffolds (KB871706, KB871620, KB871713, and KB871833). These four scaffolds map to an 8Mb region on *Astyanax* chromosome 7 (Table S3). Within this 8Mb region on Chr. 7 resides 82 annotated genes. There of the genetic markers map to the same scaffold (KB871620) and to a ∼1Mb region on Chr. 7, wherein 24 annotated genes reside.

### Sequence Analysis

A population genomic analysis was performed using DNA from wild-caught specimens from Río Gallinas (surface fish Rascón population), Río Choy (surface fish Choy population from which laboratory surface fish are derived), as well as Pachón, Tinaja and Molino caves in Mexico. We used a n=10 per population for sequence assessment. Population genomic metrics and analysis procedures are outlined in Riddle et al. 2021. cDNA sequences were aligned using SnapGene (v6.1.2), from which fixed coding sequence changes were noted (Table 1). We identified known zebrafish, mouse and human phenotypes associated with candidate genes using the BioMart tool in Ensembl (v104; Moran et al. 2022).

**Table 1.** Genetic alterations identified in candidate genes associated with bite differences.

|  |  |  |  |  |  |
| --- | --- | --- | --- | --- | --- |
| Gene | Name | Location | Genetic Alteration | Amino Acid Change | Cavefish Population affected |
| *RAB19* | *RAB19, member RAS oncogene family* | 7:8404412-8414687 | Nonsynonymous SNP | Histidine -> Glutamine | Pachón, Molino, and Tinaja |
| *arfgap3* | *ADP ribosylation factor GTPase activating protein 3* | 7:8446792-8462735 | Nonsynonymous SNP | Proline -> Histidine | Pachón |
| *pacsin2* | *Protein kinase C and casein kinase substrate in neurons protein 2* | 7:8464067-8496854 | Nonsynonymous SNP | Phenylalanine -> Leucine | Pachón and Molino |
| *large1* | *LARGE xylosyl- and glucuronyltransferase 1* | 7:9076709-9165670 | Nonsynonymous SNP | Aspartic acid -> Asparagine | Pachón, Molino, and Tinaja |
| *USP15* | *ubiquitin specific peptidase 15* | 7:9928859-9967471 | 1-13bp deletions | - | Pachón, Molino, and Tinaja |

## Results

### Cavefish exhibit differences in jaw morphology compared to surface fish

Cavefish harbor an underbite, compared to an overbite or even occlusion in surface fish, which manifests as an elongated lower jaw and a wider mouth opening or “gape” (Fig. 1 A, D). Gape was measured by taking the maximum angle of the maxillary bone to the lateral mandible. Cavefish averaged a significantly higher gape angle ranging from 130°-139° (mean= 136°), compared to a range of 96°-116° in surface fish (mean =106°; Fig. 1B). Surface x cavefish F_2_ hybrids were separated into “overbite” and “underbite” groups. F_2_ hybrids scored as having an overbite had an average gape angle of 116°, compared to an average angle of 130° in F_2_ hybrids with an underbite (Fig. 1B). An ANOVA revealed significant variation in gape angle across populations (F=18.83; p<0.001). A post hoc Tukey test showed significant differences between overbite and underbite groups at p< 0.05 (See Table S1). In addition to a larger gape angle, F_2_ hybrids with an underbite have significantly longer lower jaws (normalized mandible length) compared to overbite F_2_ hybrids (p< 0.05; Fig. 2I). No sex differences were observed for any of the jaw morphology metrics analyzed.

### A novel behavioral assay illustrates that fish with an underbite feed differently on substrate compared to fish with an overbite

Cavefish display a difference in feeding posture compared to surface fish (Schemmel, 1980; Kowalko et al. 2013). To determine if F_2_ hybrids with an underbite feed at a similar angle to cavefish (and if hybrids with an overbite feed like surface fish), we co-opted the feeding behavior assay used by Kowalko et al. 2013. Consistent with previous findings, cavefish fed at the expected posture with an average of 54°, compared to surface fish that fed at an average angle of 80° (Fig. 1C; 2A-B, Fig. S1 A-B). F_2_ hybrids with an overbite displayed a similar feeding angle to surface fish, with individual trial averages ranging from 76°-81° (post hoc Tukey p>0.05; Fig. 2C).

Surprisingly, F_2_ hybrids with an underbite did not recapitulate cavefish feeding posture, feeding at a wider range of 62°-90° (Fig. 1C; 2D). An ANOVA revealed significant variation in feeding angle across populations (F=19.01; p<0.001). Further, feeding angle is negatively correlated with gape angle (R= -0.4134; Fig. 1F), suggesting that fish with a larger gape feed at more acute angles.

Despite differing from cavefish feeding posture, F_2_ hybrids with an underbite do display interesting feeding behaviors. Compared to hybrids with an overbite (90°), F_2_ hybrids with an underbite had a maximum feeding behavior of 110°, extending their lower jaws and feeding at an almost upside-down posture (p<0.001; Fig. 1E; Fig. S1E).

Cavefish exhibit an increase in the number of teeth on both the upper and lower jaws (Atukorala et al. 2013). Tooth number was counted for F_2_ hybrids post behavior assay. There was no significant difference in tooth number in the upper jaw between overbite and underbite hybrids (p>0.05; Fig. 2J). However, F_2_ hybrids with an underbite have significantly more teeth in their lower jaw compared to F_2_ hybrids with an overbite (p<0.001; Fig. 2K). Taken together, F_2_ hybrids with an underbite have similar morphology to cavefish, with an elongated lower jaw and an increase in tooth number.

We were able to take a closer look at feeding behavior by designing a method for extracting bite impressions during behavior trials. Food carpets (see Methods) were used to visualize the number of tooth marks made by a fish during a feeding strike (Fig. 2E-H). Surface fish, feeding at ∼90°, were observed using their upper jaw to bite into the food carpet, leaving smaller bite impressions with an average of 6.7 tooth marks (Fig. 2E, L). In contrast, cavefish were observed using their lower jaws to make larger bite impressions, averaging 10 tooth marks per bite (Fig. 2F, L). F_2_ hybrids with an overbite displayed similar biting behavior to surface fish, averaging 5.6 tooth marks per bite (Fig. 2G, L). Like cavefish, F_2_ hybrids with an underbite made large bite impressions, averaging 8.6 tooth marks per bite (Fig. 2H, L). An ANOVA revealed significant variation in gape angle across populations (F=10.71; p<0.001). A post hoc Tukey test showed significant differences between overbite and underbite groups at p< 0.05 (See Table S2).

### *Bite differences are under genetic control in* Astyanax

Quantitative trait loci (QTL) analysis was performed to assess whether bite differences in cavefish are associated with genetic loci. Within our F_2_ mapping pedigree, ∼25% of individuals were scored as having an underbite (Fig. 3B). A significant QTL peak was recovered for the bite phenotype that rose above the significance threshold (p<0.05 LOD is 4.01) with a LOD score of 4.708 on linkage group (LG) 1 (Fig. 3C-D). The percent variance (PVE) explained by the bite phenotype is 9.4%. Seven genetic markers reside under the QTL peak with LOD scores ranging from 4.032 to 4.708 along a ∼5cM region on linkage group 1 (Table S3). The phenotypic effect for flanking genetic markers revealed that the homozygous cavefish genotype is associated with the underbite phenotype, while the heterozygous and homozygous surface fish genotypes are associated with an overbite (Fig. 3E).

A ∼9cM region at the end of LG 1 was anchored to four Pachón cavefish annotated genome scaffolds (AstMex102; McGaugh et al. 2014; Table S3). The analogous scaffold regions mapped to an ∼8Mb region of chromosome (Chr.) 7 on the surface fish genome (Warren et al. 2021; Fig. 4). A list of 84 annotated genes was assembled from the interval of 4 to 12 Mb on Chr. 7 (Fig.4; Table S3).

### Candidate genes for bite differences exhibit genetic alterations

To determine if cavefish harbor genetic alterations in candidate genes within the QTL interval, genomic sequences from wild-caught fish from multiple populations were assessed. We discovered three genes that had sequence alterations in Pachón cavefish, and also in two other populations (Molino and Tinaja; Table 1). The gene *RAB19*, a member of the RAS oncogene family, is predicted to have a nonsynonymous single nucleotide polymorphism resulting in a single amino acid substitution (H164G) in all three cavefish populations compared to cDNA sequences in both Rascón and Choy surface fish populations. Next, we discovered a predicted single amino acid substitution (P412H) only present in the Pachón population for the gene *arfgap3*, known as ADP ribosylation factor GTPase activating protein 3, compared to surface fish.

Three genes with identified genetic alterations have known roles in bone development and homeostasis. A single amino acid substitution (F310L) was predicted for the gene *pacsin2*, known as protein kinase C and casein kinase substrate in neurons protein 2, in the Pachón and Molino populations compared to surface fish. Based on annotations extracted from BioMart, alterations in the *pacsin2* gene result in abnormal bone mineralization. Next, a single amino acid substitution (D721N) was predicted for the gene LARGE1, known as large xylosyl- and glucuronyltransferase 1 in all three cavefish populations. Annotations for the LARGE1 gene suggest that mutations result in abnormal tongue morphology and bone structure. Finally, we discovered a putative deletion, ranging from 1-13 base pairs (potentially impacting amino acid positions 413-417) depending on the individual cavefish and population, in the gene USP*15*, known as ubiquitin specific peptidase 15. Alterations to *USP15* result in increased bone mineral density. Further, *USP15* has been shown to enhance bone morphogenetic protein signaling by targeting ALK3/BMPR1A (Herhaus et al. 2014). While it is presently unclear how these alternations impact jaw growth in cavefish, these are candidates worth pursuing in future studies.

## Discussion

Bite morphology is of functional importance for feeding, communicating, and breathing. As humans evolved smaller jaws, issues of malocclusion, tooth crowding, and facial pain arose (Kahn et al. 2020). Despite the increased in frequency of these aberrations, the precise genetic mechanisms controlling jaw size remain unclear. Here, we capitalize on the natural variation of jaw size and bite differences in divergent forms of teleost fish. We discovered that bite differences are indeed under genetic control in cavefish.

While the majority of previously studied craniofacial traits appear to be under complex genetic control in cavefish (Gross et al. 2014), we discovered a single QTL peak for the bite phenotype, with the frequency near a 3:1 (overbite: underbite) ratio in the F_2_ pedigree, suggesting a Mendelian pattern of inheritance with the surface fish alleles being dominant. However, the QTL explains <10% of the variance so it is possible that this is unlikely a monogenic trait and multiple genes or networks may be impacted. Five of the genes in the QTL region (*RAB19, arfgap3, pacsin2, LARGE1*, and *USP15*) exhibit fixed nonsynonymous mutations in cavefish compared to surface fish.

We found that some mutations (*RAB19, LARGE1*, and *USP15*) were present in all three populations of the cavefish (Pachón, Molino and Tinaja) we investigated. However, other mutations were only present in one or two populations compared to the surface fish. A potential explanation for this is that different cavefish populations may employ different genetic mechanisms to converge on similar phenotypes. An example of this is a mutation in the insulin receptor (*insra*) governing glucose intolerance in Pachón and Tinaja populations, but not Molino (Riddle & Aspiras et al. 2018). While the genes *pacsin2, LARGE1*, and *USP15* have been previously implicated in altered bone mineralization, none of the candidates have been specifically linked to changes in jaw morphology and may play novel roles in controlling bone size differences. Future functional analysis studies are needed to uncover the precise role of these genes in jaw development.

It is also possible that bite differences may not be mediated solely by a genetic mutation affecting the amino acid sequence, but rather a change in temporal or spatial gene expression during development. Protruding lower jaws have been observed in larval cavefish (Jeffrey, 2001; Yamamoto et al. 2009), suggesting that lower jaw cartilage (Meckel’s cartilage) may lay down the foundation for jaw size differences observed in adult skulls. Bone morphogenetic protein (BMP) signaling is a key regulator of endochondral ossification and has been shown to stimulate cell differentiation during cartilage development (Kobayashi et al. 2005). Allelic variation and expression of *bmp4* have been implicated in differences in cichlid jaw shape (Albertson & Kocher, 2006).

One candidate gene exhibiting sequence deletions in cavefish, *USP15*, is a known regulator of BMP signaling and may play a role in chondrogenesis of the jaw (Herhaus et al. 2014). Another gene within the QTL region is *wnt7ba*, which together with ortholog *wnt7bb* are expressed in the developing zebrafish head as early as 24 hours post fertilization (Duncan et al. 2015) and wnt/beta-catenin signaling has been shown to induce cartilaginous matrix remodeling (Yuasa & Iwamoto, 2006). Together, *USP15* and *Wnt7ba* should be further investigated across jaw development to determine if changes in expression result in an increase in lower jaw cartilage in cavefish.

While jaw size and dentition differences have been previously characterized in cavefish, the evolutionary mechanism underlying these changes remains unclear. Varying degrees of eye degeneration, shown through lens ablation studies, does not affect the length of the lower jaw (Dufton et al. 2012) and we observed no overlapping QTL for eye size, nor did we find any correlations with eye size and any jaw metrics presented here. Kowalko et al. (2013) determined that cavefish feed at a more acute angle compared to surface fish, but we did not find that F_2_ hybrids exhibiting an underbite feed at the same posture as cavefish. Further, multiple QTL for feeding angle were discovered (Kowalko et al. 2013), but do not overlap with the bite phenotype. This suggests that feeding angle is controlled by a different genetic mechanism than jaw morphology in cavefish. A previously discovered QTL for jaw angle (ventral jaw width) does map to chromosome 7, but not at the same genomic position (7:29,668,292-29,697,069) and a different scaffold (KB871834.1:596.975) than the bite QTL. Another previously characterized QTL for lower jaw size (Protas et al. 2007) maps to Chr. 14 near the gene *ghrb* (Berning et al. 2019). From these studies we can infer that the size of the adult lower jaw is likely controlled by different loci than lower jaw protrusion or bite. Besides bone and cartilage, other features within the cranium may contribute to bite differences, such as potential muscle or joint differences.

While bite differences do not correlate with feeding angle, we did discover that an underbite is associated with differences in feeding strategy, such that fish with an underbite used their lower jaws, exposing more teeth in each strike compared to fish with an overbite. This is consistent with findings in cichlids, wherein fish with shorter, stout jaws feed on hard substrate, while fish with elongated jaws can range from suction feeders to predators (Albertson et al. 2005). Further, fish exhibiting a short dentary, with long distances between the quadrate joint and opening/closing ligaments feed on attached foods, such as algae and microinvertebrates, requiring a greater force to remove from surfaces (Husley et al. 2010). This is consistent with what surface fish likely encounter in terms of feeding ecology.

In the caves, however, there is no photosynthetic input and few available prey. Why then would cavefish need to evolve wider, longer jaws with more teeth? Espinasa et al. (2017) analyzed gut contents from wild-caught cavefish from the Pachón cave during both the rainy and dry seasons and determined adult cavefish mainly subside on a diet of bat guano and detritus. This suggests that cavefish use their larger jaws and increased tooth number to filter feed through the muddy cave pool floor. Additionally, cavefish have an increase in tastebud number, both extraorally and specifically within the lower jaw extending along the lingual epithelium toward the posterior part of the jaw (Varatharasan et al. 2009). Cavefish may have evolved an increase in jaw size and wider gape to expose more tastebuds, thus increasing taste sensitivity in a nutrient poor environment. Alternatively, tooth and jaw differences may have evolved as a consequence of indirect selection (Jeffery, 2010), wherein sensory enhancements such as increased cranial innervation (Sumi et al. 2015) and taste bud number were under selection, causing pleiotropic changes that resulted cranial modifications. Further studies using genetic perturbations will uncover the precise mechanisms governing these changes. Taken together, we have established cavefish as a powerful genetic model for understanding evolutionary changes in morphology and behavior, particularly in the context of jaw evolution.

## Figure Legends

**Supplemental Figure 1. Ethograms illustrate feeding posture differences between surface, cavefish and hybrids**. Consistent with findings from Kowalko et al. 2013, we determined that surface fish have a near perpendicular feeding posture with an average angle between 80°-90° (A) and cavefish feed at a lower angle of 40°-60° (B). Surface x Pachón F_2_ hybrids demonstrated three feeding posture categories: surface-like F_2_ hybrids with an average feeding angle between 80°-90° (C), a mix of surface- and cave-like feeding postures with angles ranging from 40°-90° (D), and an extreme obtuse posture with angles up to 110° (E).

