## Supplementary figures and images for "Genetic mapping of craniofacial traits in the Mexican tetra reveals loci associated with bite differences between cave and surface fish"

### Supplemental Figure 1

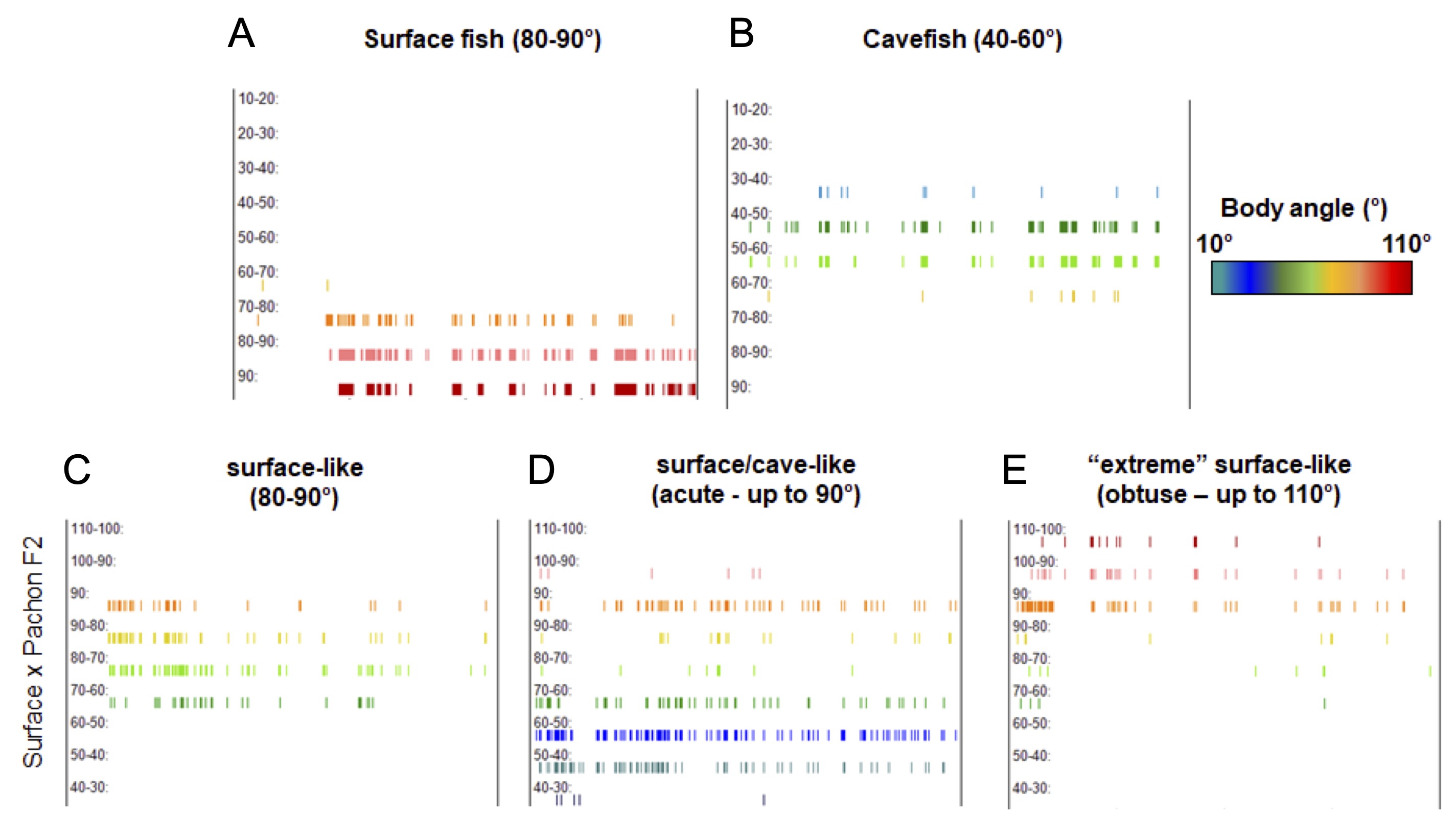

### Supplemental Table 1

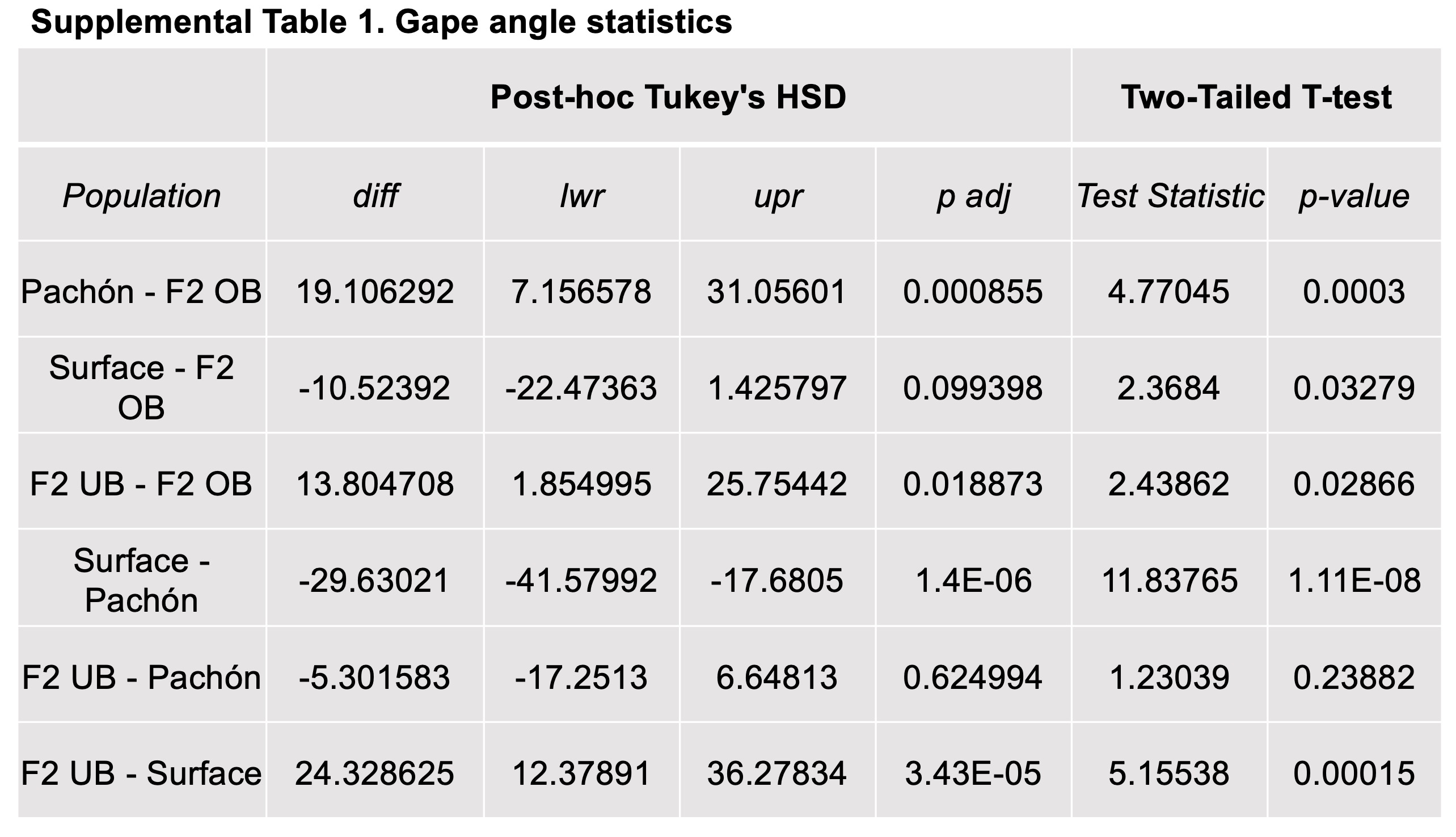

### Supplemental Table 2

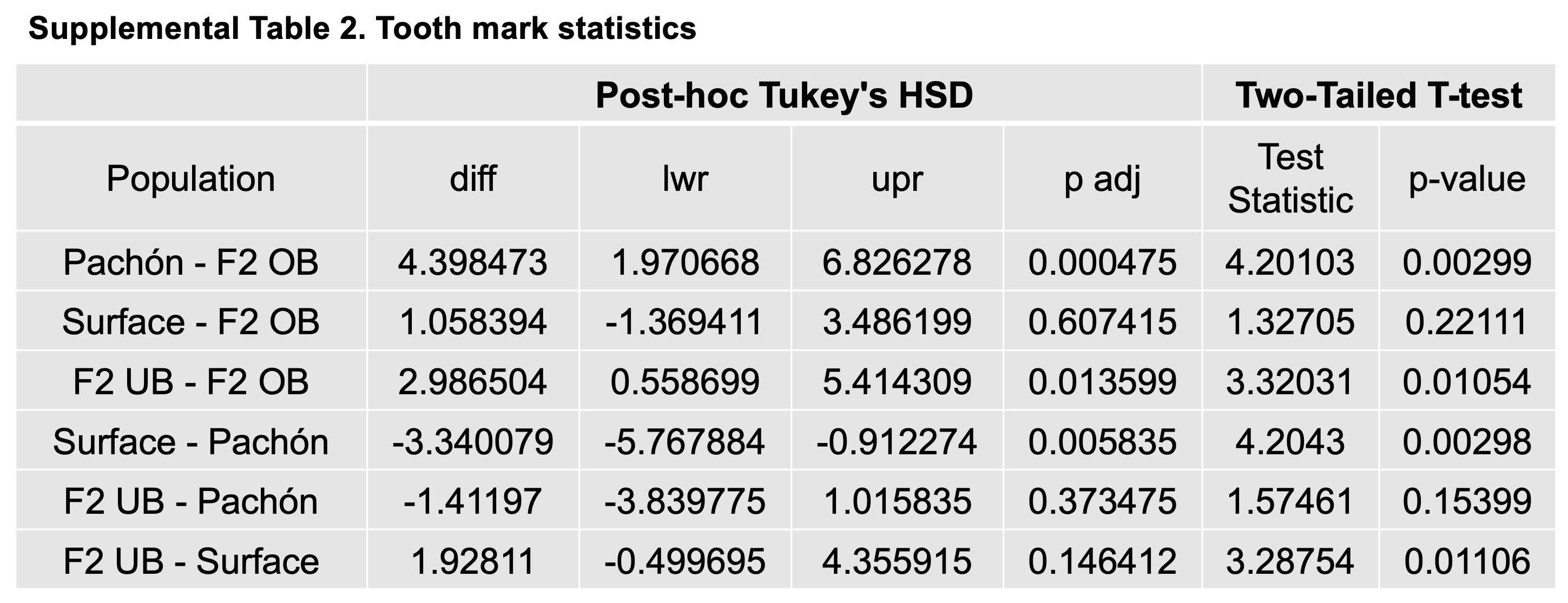

### Supplemental Table 3

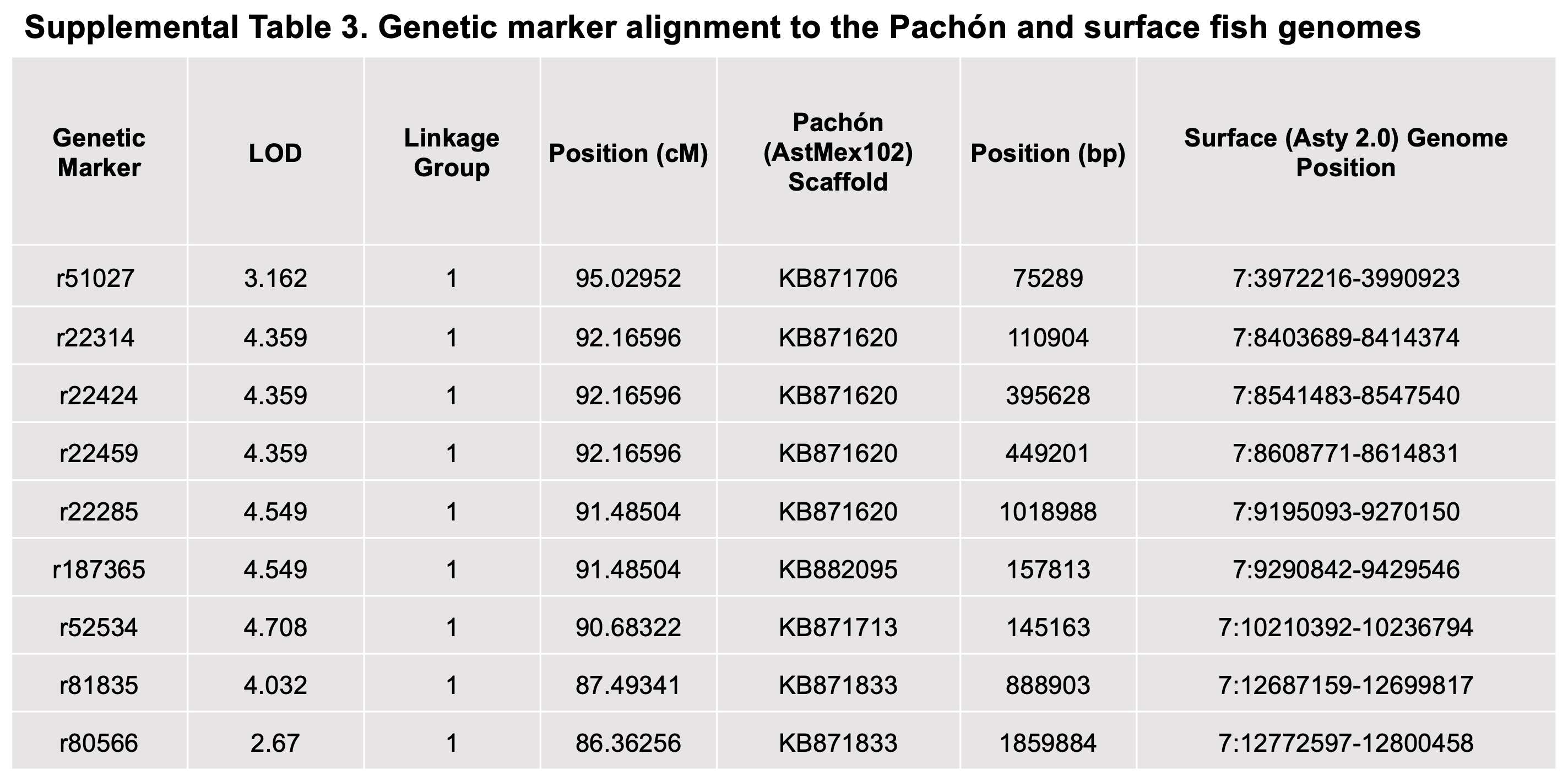
